# Native Proteomics by Capillary Zone Electrophoresis-Mass Spectrometry

**DOI:** 10.1101/2024.04.24.590970

**Authors:** Qianyi Wang, Qianjie Wang, Zihao Qi, William Moeller, Vicki H Wysocki, Liangliang Sun

**Author notes:** Those three authors contributed equally to this work (Qianyi Wang, Qianjie Wang, Zihao Qi).

## Abstract

Native proteomics measures endogenous proteoforms and protein complexes under a near physiological condition using native mass spectrometry (nMS) coupled with liquid-phase separations. Native proteomics should provide the most accurate bird’s-eye view of proteome dynamics within cells, which is fundamental for understanding almost all biological processes. nMS has been widely employed to characterize well-purified protein complexes. However, there are only very few trials of utilizing nMS to measure proteoforms and protein complexes in a complex sample (i.e., a whole cell lysate). Here, we pioneer the native proteomics measurement of large proteoforms or protein complexes up to 400 kDa from a complex proteome via online coupling of native capillary zone electrophoresis (nCZE) to an ultra-high mass range (UHMR) Orbitrap mass spectrometer. The nCZE-MS technique enabled the measurement of a 115-kDa standard protein complex while consuming only about 0.1 ng of protein material. nCZE-MS analysis of an *E*.*coli* cell lysate detected 72 proteoforms or protein complexes in a mass range of 30-400 kDa in a single run while consuming only 50-ng protein material. The mass distribution of detected proteoforms or protein complexes agreed well with that from mass photometry measurement. This work represents a technical breakthrough in native proteomics for measuring complex proteomes.

---

Proteins regulate cellular processes by their diverse proteoforms^[1,2]^ and the various protein complexes via non-covalent protein-protein interactions, protein-ligand bindings, and protein-DNA/RNA interactions.^[3]^ Native mass spectrometry (nMS) provides essential insights into the structures, functions, and dynamics of proteoforms and protein complexes near physiological conditions.^[4–7]^ nMS has been widely employed to study well-purified proteoforms and protein complexes with low complexity through either direct infusion^[8-13]^ or coupling with online/offline native separation methods, including size-exclusion chromatography (SEC),^[14-18]^ ion-exchange chromatography (IEX),^[19,20]^ hydrophobic interaction chromatography (HIC),^[21,22]^ and capillary zone electrophoresis (CZE).^[23]^ Native proteomics aims to measure endogenous proteoforms and protein complexes under a near physiological condition on a proteome scale and it requires highly efficient separation techniques for protein complexes prior to nMS.^[24]^ The first native proteomics study coupled off-line IEX or native gel-eluted liquid fractionation with direct infusion nMS for the characterization of protein complexes in mouse heart an human cancer cell lines, identifying 125 endogenous complexes from about 600 fractions.^[25]^ More recently, direct infusion nMS was employed to measure protein complexes from a human heart tissue lysate using a Fourier-transform ion cyclotron resonance (FTICR) mass spectrometer with the identification of a handful of protein complexes about 30 kDa or smaller.^[26]^ Native CZE-MS (nCZE-MS) has high separation efficiency and high detection sensitivity for protein complexes and has been applied to analyzing low-complexity protein samples, i.e., monoclonal antibodies,^[27]^ large protein complexes like GroEL (near 1MDa),^[28-30]^ ribosomes,^[31]^ and nucleosomes.^[32]^ Native SEC fractionation and online nCZE-MS analysis of an *E. coli* cell lysate identified 23 protein complexes smaller than 30 kDa, representing the first native proteomics study of a complex proteome using online liquid-phase separation-MS.^[33]^ However, those native proteomics studies are either too time and labor-consuming or only able to detect small proteoforms/protein complexes from complex proteomes.

In this study, we developed a high-throughput nCZE-MS technique for native proteomics measurement of large proteoforms and protein complexes up to 400 kDa from complex samples, *i*.*e*., an *E. coli* cell lysate. The nCZE-MS technique is based on the online coupling of nCZE to an ultra-high mass range (UHMR) Orbitrap mass spectrometer. We first evaluated the nCZE-MS technique using a standard protein complex mixture. Then, we employed the technique to analyse endogenous proteoforms and protein complexes in *E. coli* cells. We also compared our nCZE-MS data with mass photometry results in terms of the mass distribution of *E. coli* proteoforms and protein complexes.^[34]^

Figure 1. shows the workflow of native proteomics analysis of an *E. coli* cell lysate using our nCZE-UHMR Orbitrap platform. Briefly, the cultured *E. coli* cells (Top10 strain) were lysed in a Dulbecco’s phosphate-buffered saline (dPBS) buffer containing complete protease inhibitors and phosphatase inhibitors. The cell lysate was then buffer-exchanged on a spin column (Bio-Rad P6) to a buffer containing 20 mM ammonium acetate (AmAc, pH ∼7.0) by gel filtration, followed by nCZE-MS analysis. The online nCZE-MS was assembled by coupling a Sciex CESI-8000 Plus capillary electrophoresis (CE) autosampler to a Thermo Fisher Scientific Q-Exactive UHMR mass spectrometer through a commercialized electrokinetically pumped sheath flow CE-MS interface (EMASS-II, CMP Scientific).^[35,36]^ A 1-meter-long linear polyacrylamide (LPA) coated capillary (50-μm i.d., 360-μm o.d.) was used for the CZE separation, and the LPA coating was employed to reduce the protein non-specific adsorption onto the capillary inner wall. The background electrolyte (BGE) for CZE was 25 mM AmAc (pH ∼7.0), and the sheath buffer for electrospray ionization (ESI) was 10 mM AmAc (pH ∼7.0). Only roughly 50 ng of the *E. coli* sample was consumed in a single nCZE-MS run. Raw MS data were averaged every 30 seconds, followed by mass deconvolution using UniDec and ESIprot.^[37, 38]^ The detailed experimental procedure is described in the **Supporting Information**.

**Figure 1.**
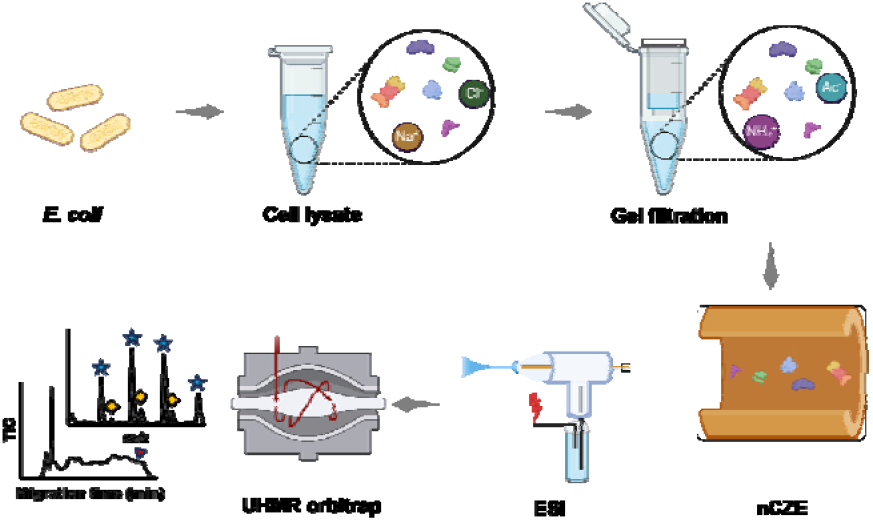
Flow chart of nCZE-ESI-MS for native proteomics of an *E. coli* cell lysate. The figure is created using the BioRender and used here with permission.

We investigated the sensitivity of the nCZE-ESI-UHMR platform for measuring protein complexes using a mixture of standard proteins and protein complexes, **Figure S1**. High intensity was observed for streptavidin (SA, 53 kDa), carbonic anhydrase (CA, 29 kDa), C-reactive protein (CRP, 115 kDa), and bovine serum albumin (BSA, 66 kDa) in the original sample via only consuming about 15 ng of those proteins. After sample dilution by a factor of 50, a clear CRP peak was still observed, even though only 0.1 ng of the protein complex was loaded, indicating the high sensitivity of the technique. **Figure S2A** shows one mass spectrum of three SA tetramers with masses of 53084.67 Da, 53216.07 Da, and 53347.97 Da. A 131-Da mass difference was observed between neighboring SA complexes, corresponding to N-terminal methionine variation on SA, which is consistent with the literature.^[39]^ **Figure S2B** shows a mass spectrum of the CA-Zn(II) complex (29088.10 Da) and another CA complex (29194.01 Da) with an additional 107-Da mass shift compared to the CA-Zn(II) complex.^[39,40]^ **Figure S2C** shows the mass spectrum of the pentameric CRP complex in the original sample. Based on De La Mora’s prediction of the maximum (Rayleigh) charge ‘Z_R_’ of a native protein during the ESI process (Z_R_ = 0.0778*M^0.5^), the max charge of CRP is around 26.4.^[41,42]^ The max charge states of CRP observed in the original and 50-time diluted samples are 27 and 26, matching well with the ZR of native CRP. We observed slightly lower max charge states compared to the theoretical charge states for the SA tetramer, CA-Zn (II) complex, and BSA, **Figure S2D**. The data clearly demonstrate that intact protein complexes are maintained in native-like states during nCZE-ESI-UHMR measurements.

The high sensitivity of nCZE-UHMR for the standard protein complexes motivated us to analyze an *E. coli* cell lysate. **Figure 2A** shows an example electropherogram of the sample from nCZE-MS. The proteoforms or protein complexes migrated out of the capillary in a time range of 20-65 minutes, allowing the mass spectrometer sufficient time for data acquisition (i.e., acquiring mass spectra and tandem mass spectra). In total, we detected 99 proteoforms or protein complexes in a mass range of 10-400 kDa after spectrum averaging and mass deconvolution. Information on the detected proteoforms or protein complexes is listed in **Table S1. Figures 2B-2F** show the mass spectra of some examples larger than 40 kDa, i.e., ∼41, 139, 146, 318, 340, and 387 kDa. Those proteoforms or protein complexes show native-like and clear mass spectra. For example, **Figure 2E** shows two co-migrating proteoforms or protein complexes with masses ∼318 and ∼340 kDa. Their most-abundance charge states are +34 and +36, respectively. The largest proteoform or protein complex detected in this study is ∼387 kDa, carrying around 42 charges (**Figure 2F**). Some additional examples are shown in **Figure S3**.

**Figure 2.**
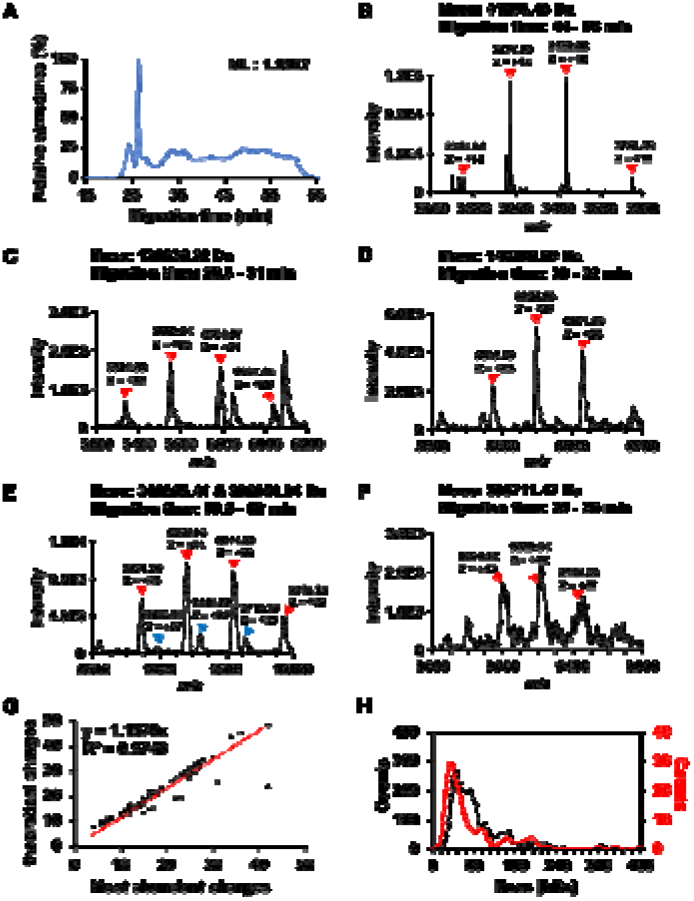
Summary of detected proteoforms or protein complexes from an *E. coli* cell lysate using nCZE-ESI-UHMR. (A) Representative electropherogram of nCZE-ESI-UHMR analyses of the *E. coli* cell lysate. (B)-(F) Mass spectra of five examples of large proteoforms/protein complexes detected. The charge states and deconvolved mass of each proteoform/protein complex are labelled. (G) Linear correlation between the most abundant charges and theoretical Rayleigh charges (Z_R_) of all proteoforms/protein complexes detected in single-shot nCZE-UHMR. (H) Alignment of the mass distribution of proteoforms/protein complexes in the *E. coli* cell lysate from mass photometry (black dash line) and nCZE-UHMR (red line) analyses.

We further examined the correlation between the predicted Rayleigh charge (Z_R_) from De La Mora’s theory and the experimental maximum charge state of detected proteoforms or protein complexes, **Figure 2G**. ^[41,42]^ We used the most abundant charge state instead of the highest charge state for each proteoform/protein complex here to avoid potential variations introduced during manual determination of the highest charge state. We observed a strong linear correlation (R^2^ = 0.97, slope of 1.16) between the experimental and predicted charge states. The slope indicates that the theoretical charges are slightly higher than the most abundant charges, suggesting the preservation of native states of the proteoforms or protein complexes in this experiment. We further employed mass photometry (MP) to collectively obtain the rough mass distribution for proteoforms/protein complexes in the same *E. coli* cell lysate in a nearly physiological solution based on the quantification by light scattering.^[34,43,44]^ The masses of proteoforms/protein complexes range from 10 kDa to 400 kDa according to the MP data, **Figure 2H** (black dashed line). About 72% of the molecule counts (2558 of 3555) from the MP analysis are smaller than 100 kDa. Interestingly, the molecular mass distributions from the MP and nCZE-MS analysis agree reasonably well, **Figure 2H**, considering the low mass cutoff of MP. For example, the largest proteoform or protein complex detected by nCZE-MS is close to 400 kDa, and 78% (77 out of 99) of the proteoforms/protein complexes from nCZE-MS are smaller than 100 kDa, **Table S1**. It has been demonstrated that nMS and MP can produce consistent mass assessments of large proteins or protein complexes and offer complementary information about the analytes.^[45]^

Our native proteomics study here is important because, for the first time, we can achieve a proteome-scale measurement of endogenous proteoforms and protein complexes in a complex biological sample under near-physiological conditions by nMS with relatively high throughput. Nearly 100 endogenous intact proteoforms and protein complexes up to 400 kDa were detected from an *E. coli* cell lysate by online nCZE-MS in roughly 1-hour measurements with the consumption of 50-ng protein material. nCZE-MS can maintain the protein molecules from a complex cell lysate in close-to-native states during the measurement, evidenced by the strong linear correlation between the predicted Rayleigh charge ‘Z_R_’ and experimental most-abundance charge state of detected proteoforms or protein complexes, as well as the strong agreement in molecular mass distributions between the nCZE-MS and MP data.

Compared with native SEC-MS, another well-recognized technique for native proteomics, nCZE-MS has better sensitivity due to higher separation resolution and a much lower flow rate for ESI. However, native SEC-MS is robust and has high throughout. ^[15,46]^ We expect that coupling of native SEC fractionation with nCZE-MS will be helpful for further boosting the proteome coverage of native proteomics because the two separation techniques offer orthogonal separations of protein complexes.

The current study still has several limitations. Firstly, we only observed the mass information of proteoforms or protein complexes and did not generate high-quality MS/MS data during the nCZE-MS run, impeding the accurate identification of each protein. Those detected proteoforms belong to level 5 identifications.^[47]^ We will solve this issue by optimizing surface-induced dissociation (SID) or higher energy collisional dissociation (HCD) to achieve better fragmentation of large proteoforms or protein complexes in our future study. Second, the sample loading capacity of nCZE is low, impeding the detection of low-abundance proteoforms or protein complexes and reducing the quality of acquired MS/MS spectra. We will enhance the overall sample loading capacity of nCZE by some online stacking techniques (e.g., capillary isoelectric focusing ^[27]^) or offline fractionation techniques (e.g., SEC ^[33]^). Third, the separations of large protein complexes by nCZE need to be further improved regarding separation peak capacity an reproducibility. **Figure S4** shows the electropherograms of triplicate nCZE-MS measurements of the *E. coli* cell lysate. **Figure S5** shows the extracted ion electropherograms of two example proteoforms/protein complexes. The peaks are much wider than those for denaturing CZE. The roughly estimated peak capacity of the nCZE separation is 15 based on the separation window and the average full peak width at half maximum of the two examples in **Figure S5**. The relatively low peak capacity is possibly due to the protein dispersion under the applied pressure and non-specific protein adsorption on the capillary inner wall. The separation profiles have some significant changes after 45 min in the second and third runs compared to the first run, most likely due to changes at the capillary inner wall after the first run of the *E. coli* sample. We need to develop procedures to clean up the capillary inner wall between nCZE-MS runs^[48]^ and improve the capillary inner wall coating through different chemistries, *e*.*g*., carbohydrate-based neutral coating,^[27]^ to reduce protein adsorption for better separation peak capacity and reproducibility. Lastly, the bioinformatics tool for data analysis needs to be improved. We employed mass deconvolution using UniDec ^[37]^ and ESIprot ^[38]^ for each averaged mass spectrum across the whole run. This approach was tedious and could be problematic for low-abundance proteoforms or protein complexes. More efforts ar needed to build streamlined bioinformatic tools for large-scale native proteomics using, e.g., nCZE-MS.

In summary, we have demonstrated, for the first time, that nCZE coupled to an Orbitrap UHMR mass spectrometer is an effective and sensitive platform to measure large proteoforms or protein complexes up to 400 kDa from a complex proteome sample. This nCZE-MS technique enabled highly sensitive detection of standard protein complexes via consuming only pg amounts of protein material. The technique successfully detected nearly one hundred proteoforms or protein complexes from an *E. coli* cell lysate in a mass range of 10-400 kDa. With further improvements in gas-phase fragmentation and nCZE separation peak capacity and reproducibility, we envision that nCZE-orbitrap UHMR will become a powerful tool in native proteomics of complex proteome samples.

## Supporting Information

The authors have cited additional references within the Supporting Information.^[49-51]^

## Author Contributions

Qianjie Wang prepared the *E. coli* cell lysate and did the sample preparation. Qianyi Wang did the data analysis. Qianyi Wang and Qianjie Wang wrote the draft of the manuscript. Qianyi Wang, Qianjie Wang, Zihao Qi, and William Moeller worked together for the native CZE-MS system setup and analysis of the prepared *E. coli* cell lysate. Zihao Qi generated the mass photometry data. Vicki H Wysocki and Liangliang Sun designed the experiments, oversaw the project, and edited the manuscript. All authors made comments on the manuscript.

## Acknowledgements

The authors thank the support from the National Cancer Institute (NCI) through grant R01CA247863 (Sun), the National Institute of General Medical Sciences (NIGMS), through grants R01GM125991 (Sun) and R01GM118470 (Sun), and the National Science Foundation through the grant DBI1846913 (CAREER Award, Sun). This research was supported by NIH Native Mass Spectrometry-Guided Structural Biology Center (RM1GM149374 to V.H.W.)

## Entry for the Table of Contents

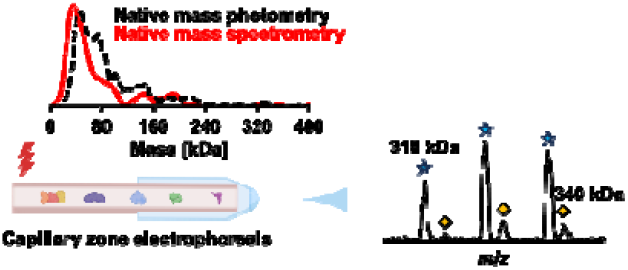

Native proteomics enabled the measurement of protein complexes up to 400 kDa from a complex proteome, via online coupling of native capillary zone electrophoresis to an ultra-high mass range Orbitrap mass spectrometer. Mass spectrometry-based native proteomics agreed well with native mass photometry regarding the mass distribution of detected proteins.

