## supporting information I for "Native Proteomics by Capillary Zone Electrophoresis-Mass Spectrometry"

* Corresponding author.

| **Content** | **Page** |
| --- | --- |
| Experimental Procedures | S3-S5 |
| Table S1 | S6-S8 |
| Figure S1 | S9 |
| Figure S2 | S10 |
| Figure S3 | S11 |
| Figure S4 | S12 |
| Figure S5 | S13 |

**Experimental procedures**

**Materials and chemicals**

Bare fused silica capillaries (50-μm i.d., 360-μm o.d.) were purchased from Polymicro Technologies (Phoenix, AZ). 3-(Trimethoxysilyl) propyl methacrylate, ammonium persulfate, ammonium acetate (AmAc), Dulbecco’s phosphate-buffered saline (dPBS), bovine serum albumin (BSA) and carbonic anhydrase (CA) from bovine erythrocytes, Cytochrome C (Cyt C), myoglobin from equine (Myo), C-reactive protein (CRP), glutamate dehydrogenase (GDH) were purchased from Sigma-Aldrich (St. Louis, MO). Hydrofluoric acid (HF), streptavidin (SA) and LC/MS grade water were purchased from Fisher Scientific (Pittsburgh, PA). Acrylamide were purchased from Acros Organics (NJ, USA). Micro Bio-Spin^TM^ 6 kDa gel-filtration column units for buffer exchange was purchased from Bio-Rad. Protease inhibitors (cOmplete ULTRA Tables) and phosphatase inhibitors (PhosSTOP) were from Roche.

**Sample preparation**

A mixture of standard protein complexes containing Cyt C (0.7 μM), Myo (0.5 μM), CA (3 μM), SA (1.1 μM), BSA (0.7 μM), CRP (1 μM) and GDH (6 μM) was prepared in 20 mM AmAc (pH ~ 7.0).

*E. coli* (strain Top10) was cultured in Terrific Broth (TB) medium at 37 °C until OD_600_ reached 0.7. After washed with dPBS three times, 2 g pellet was suspended in 5 mL dPBS buffer plus complete protease inhibitors and phosphatase inhibitors and homogenized for 30 s, followed by sonication with a Branson Sonifier 250 (VWR Scientific, Batavia, IL) on ice for 2 minutes, 3 times. After centrifugation at 10,000 g for 10 minutes, the supernatant containing the extracted proteins was collected. A small aliquot of the diluted sample was used for the bicinchoninic acid (BCA) assay to determine the protein concentration (~2 mg/mL). One aliquot of the E. coli lysate was diluted 8,000 times (~2.5 nM assuming an average molecular weight of 80 kDa) by 20 mM AmAc and directly measured using mass photometry.

Another aliquoted *E. coli* lysate was buffer exchanged to 20 mM AmAc by Bio-Spin 6 kDa gel-filtration column. The column which was washed with 20 mM AmAc and centrifuged at 1,000 x (g) for 2 minutes and repeated for 3 times. A 50 µL (100 µg protein) cell lysate was loaded on a 6 kDa gel-filtration and centrifuged for 4 minutes at 1,000 x (g). And the step was repeated with another pre-washed gel-filtration column to ensure the depletion of dPBS.

**Mass photometry**

Mass photometry experiments were conducted on a TwoMP instrument (Refeyn Inc.) Glass coverslips and silicone gaskets used in this measurement were cleaned with ultrapure water and isopropanol (IPA) sequentially in order of water - IPA - water - IPA - water, then dried by pure nitrogen. The oil immersion objective was covered with a clean coverslip, and a 6-well silicone gasket was placed on the top of the coverslip.

Calibration was carried out using a mixture of 10 nM thyroglobulin (TG) and beta-amylase (BAM). Four peaks corresponding to the monomer, dimer tetramer of BAM, together with the dimer of TG, have been detected. These contrasts and corresponding masses generated a calibration curve with an R square value of 0.99999, and the calibration was used to identify the rough mass of individual proteins or protein complexes existing in the cell lysate.

The vendor-reported parameters related to the mass photometry are as follows. The mass precision is 2%. The mass error is 5%. The resolution (defined as FWHM) is 25 kDa @ 66 kDa and 60 kDa @ 660 kDa, respectively.

**Preparation of LPA-coated separation capillary**

The inner wall of the separation capillary (50-μm i.d., 360-μm o.d.) was coated with linear polyacrylamide (LPA) based on the protocol described in previous references.^[49, 50]^ Briefly, a bare fused silica capillary was successively flushed with 1 M sodium hydroxide, water, 1 M hydrochloric acid, water, and methanol, followed by treatment with 3-(trimethoxysilyl) propyl methacrylate for at least 24 hours to introduce carbon-carbon double bonds on the inner wall of the capillary. The treated capillary was filled with degassed acrylamide solution in water (4%) containing ammonium persulfate, followed by incubation at 50 °C water bath for 55 min with both ends sealed by silica rubber. After that, the capillary was flushed with water to remove the unreacted reagents. Then one end of the LPA-coated capillary was etched with HF based on the protocol in reference ^[51]^ for 85 minutes to reduce its outer diameter to around 70 μm.

**Native CZE-ESI-MS**

A Beckman CESI8000 Plus capillary electrophoresis autosampler was used for the automated operation of capillary zone electrophoresis (CZE). A commercialized electrokinetically pumped sheath flow interface (CMP Scientific) was used to couple CZE to mass spectrometer.^[35,36]^ A Q-Exactive UHMR mass spectrometer (Thermo Fisher Scientific) were used for the experiments. The interface was directly attached to mass spectrometer. The ESI emitters of the interface were pulled from borosilicate glass capillaries (1.0 mm o.d., 0.75 mm i.d.) with a Sutter P-1000 flaming/brown micropipet puller with an orifice size ~25 μm. The sheath liquid contains 10 mM AmAc. Voltage for ESI was ~2 kV. A 1-meter LPA coated capillary (50-μm i.d. and 360-μm o.d.) was used for the CZE. The background electrolyte (BGE) for CZE was 25 mM AmAc (pH ~ 7.0).

The transfer capillary temperature was 250 ^o^C, and the S-lens RF level was 200. The number of micro scans was 5 for MS, and the in-source trapping (IST) desolvation voltage was -30V. The trapping gas flow was at 5 (UHV readback showed ~1E-11 mbar). The resolution setting for MS was 6250 (m/z 400). The AGC target was 1E6 for MS. The maximum injection time was 200 ms for MS. The mass range for MS scans was 1000-10000 m/z. The *E. coli* sample was injected into the separation capillary for CZE-MS/MS with 5-psi pressure for 9.5 s (50 nL, 2.5% of capillary volume, ~50 ng). A 70-minute CZE separation with 30 kV applied at the BGE end and 1 psi was applied at the mean time. For standard protein mixture, the separation is 45 min, and the separation is under 1.5 psi.

**Data analysis**

All the mass spectra were first averaged for a time window of every 30 s, followed by inputting the data into UniDec. Only peaks with S/N better than 10 were analyzed. Most of the settings of UniDec^[37]^ analysis were at default values except for applying the ‘Automatic m/z Peak Width’ and the ‘Suppress Artifacts’ with ‘Some’ or ‘Lots’. Next, the successive charge state distribution of proteoforms/protein complexes was manually checked to ensure the correct distribution. Finally, we calculated the mass and the standard deviation of the proteoform/protein complex by ESIProt^[38]^ based on the m/z of the successive charge states from UniDec. The m/z values used for ESIProt and the standard deviations of the deconvolved masses are listed in **Table S1**. We ensured that the data from ESIProt and UniDec agreed with each other for all the proteoforms/protein complexes in **Table S1**.

**Table S1. List of proteoforms/protein complexes detected from nCZE-MS analysis of an *E. coli* cell lysate.**

| **Number** | **Migrationtime (min)** | **Mass (Da)** | **m/z** | **m/z** | **m/z** | **m/z** | **Charge state for the most abundant m/z** | **standard deviation (Da)** |
| --- | --- | --- | --- | --- | --- | --- | --- | --- |
| **1** | 51-61 | 9737.84 | 1948.5 | **2435.44** | 3247.12 |  | **4** | **0.4483** |
| **2** | 43-44 | 11672.41 | **1946.35** | 2335.56 |  |  | **6** | **0.5002** |
| **3** | 36-37 | 15145.4 | 2164.62 | **2525.26** |  |  | **6** | **0.1608** |
| **4** | 55-58 | 16027.66 | 2004.46 | **2290.68** |  |  | **7** | **0.0618** |
| **5** | 61-62 | 17317.3 | **2165.66** | 2474.92 |  |  | **8** | **0.1184** |
| **7** | 46-49 | 17701.47 | **2213.7** | 2529.78 |  |  | **8** | **0.0938** |
| **6** | 52-60 | 17701.91 | 2529.81 | **2951.3** | 3541.48 |  | **6** | **0.3963** |
| **8** | 60-61 | 18118.92 | 2014.22 | **2265.86** | 2589.44 |  | **8** | **0.1039** |
| **9** | 28.5-38 | 18160.74 | 2019.02 | **2271.27** | 2595.01 |  | **8** | **2.3607** |
| **10** | 44-47 | 18859.82 | 2096.52 | **2358.51** |  |  | **8** | **0.2881** |
| **11** | 51-60 | 19476.28 | 2165.13 | **2435.44** |  |  | **8** | **1.1615** |
| **12** | 32-40 | 20713.85 | 2302.52 | 2590.25 | **2960.15** |  | **7** | **0.2076** |
| **13** | 61-62 | 22313.11 | 2480.3 | **2790.08** |  |  | **8** | **0.7443** |
| **14** | 23-25 | 24963.11 | 2270.37 | **2497.31** | 2774.71 |  | **10** | **0.1834** |
| **15** | 42-46 | 24973.98 | **2498.4** | 2775.9 |  |  | **10** | **0.0759** |
| **16** | 27-30 | 25081.3 | 2281.02 | **2509.09** | 2788 |  | **10** | **1.4563** |
| **17** | 28.5-30 | 25782.96 | 2344.9 | **2579.29** | 2865.81 |  | **10** | **0.2316** |
| **18** | 33-34 | 26220.08 | 2384.62 | **2623.05** |  |  | **10** | **0.4861** |
| **19** | 23-27 | 28474.4 | 2589.48 | **2848.46** | 3164.95 |  | **10** | **1.1474** |
| **20** | 27-27.5 | 28547.59 | 2596.23 | **2855.78** |  |  | **10** | **0.1961** |
| **21** | 31-32 | 28719.87 | 2612.03 | **2873.5** |  |  | **10** | **5.8019** |
| **22** | 25-25.5 | 29118.19 | 2648.14 | **2912.8** |  |  | **10** | **0.3766** |
| **23** | 50-53 | 29213.22 | **2435.42** | 2656.78 |  |  | **12** | **0.3871** |
| **24** | 46-50 | 29484.87 | 2458.09 | **2681.44** |  |  | **11** | **0.1645** |
| **25** | 23.5-26 | 29576.46 | **2689.77** | 2958.66 |  |  | **11** | **0.0972** |
| **26** | 34.5-38 | 29913.23 | **2720.41** | 2992.31 |  |  | **11** | **0.2847** |
| **27** | 38.5-40.5 | 29955.01 | **2724.18** | 2996.52 |  |  | **11** | **0.1608** |
| **28** | 36-45 | 30291.06 | 2525.25 | **2754.73** | 3030.14 |  | **11** | **0.2294** |
| **29** | 36-41 | 30291.19 | 1894.23 | **2020.39** | 2164.67 |  | **15** | **0.4177** |
| **30** | 31-32 | 30978.18 | **2817.22** | 3098.81 |  |  | **11** | **0.221** |
| **31** | 23-25 | 31227.76 | 2603.36 | **2839.89** | 3123.74 |  | **11** | **0.4544** |
| **32** | 49-56 | 31869.49 | 2656.82 | **2898.21** |  |  | **11** | **0.3695** |
| **33** | 26.5-28.5 | 32123.87 | 2678.01 | **2921.34** | 3213.4 |  | **11** | **0.1921** |
| **34** | 28.5-36 | 32336 | 2695.66 | **2940.63** | 3234.64 |  | **11** | **0.2808** |
| **35** | 31-32 | 32776.2 | 2522.26 | **2732.38** | 2980.62 |  | **12** | **0.3806** |
| **36** | 26.5-30 | 33405.11 | 2784.83 | **3037.81** | 3341.47 |  | **11** | **0.6682** |
| **37** | 53-57 | 35403.35 | **2951.29** | 3219.49 |  |  | **12** | **0.0584** |
| **38** | 41-43 | 36252.87 | **2266.56** | 2418.3 | 2590.31 |  | **16** | **5.8155** |
| **39** | 60-61 | 36305.6 | **3026.48** | 3301.51 |  |  | **12** | **0.1008** |
| **40** | 37-41 | 38516.95 | 2141.01 | **2266.46** | 2408.39 |  | **17** | **3.8128** |
| **41** | 52-56 | 38954.51 | 2783.48 | **2997.5** |  |  | **13** | **0.1503** |
| **42** | 59-60 | 40660.17 | 2905.42 | 3128.64 | **3389.3** |  | **12** | **1.3982** |
| **43** | 32-34 | 40706.16 | 3132.23 | **3393.21** |  |  | **12** | **0.38** |
| **44** | 51-61 | 40986.11 | 2928.51 | **3153.79** | 3416.6 |  | **13** | **1.039** |
| **45** | 44-56 | 41255.43 | 2947.79 | 3174.53 | **3438.95** | 3751.52 | **12** | **0.3716** |
| **46** | 32-38 | 41427.77 | 3187.77 | **3453.31** |  |  | **12** | **0.1998** |
| **47** | 47-51 | 41439.41 | **3188.6** | 3454.35 |  |  | **13** | **0.9952** |
| **48** | 40-44 | 42375.93 | **3027.75** | 3260.77 | 3532.38 |  | **14** | **1.343** |
| **49** | 44-47 | 42447.32 | 3033.01 | **3266.13** |  |  | **13** | **1.0201** |
| **50** | 39-43 | 42849.05 | 3297.09 | **3571.76** |  |  | **12** | **0.0301** |
| **51** | 62-63 | 43192.67 | 2880.42 | **3086.24** | 3323.59 |  | **14** | **1.295** |
| **52** | 36-40 | 45523.55 | 3035.98 | **3252.66** | 3502.77 |  | **14** | **0.9098** |
| **53** | 27.5-30 | 46037.81 | 3070.15 | **3289.47** |  |  | **14** | **0.9457** |
| **54** | 63-64 | 46310.85 | 3563.08 | **3860.57** |  |  | **12** | **5.5207** |
| **55** | 41-45 | 48346.97 | **3224.16** | 3454.34 |  |  | **15** | **0.4473** |
| **56** | 43-46 | 50193.31 | 3347.25 | **3586.22** |  |  | **14** | **0.4685** |
| **57** | 34-36 | 53944.84 | **3597.34** | 3854.2 |  |  | **15** | **0.2069** |
| **58** | 44-47 | 54162.8 | 3386.23 | **3611.93** | 3869.65 |  | **15** | **1.5673** |
| **59** | 48-49 | 54370.69 | 3399.16 | **3625.67** | 3884.7 |  | **15** | **0.9051** |
| **60** | 26-27 | 56947.99 | **3797.56** | 4068.7 |  |  | **15** | **0.419** |
| **61** | 54-57 | 60544.4 | 3562.39 | **3784.93** | 4037.47 |  | **16** | **2.2286** |
| **62** | 43-44 | 60584.83 | 2635.21 | **2754.78** |  |  | **22** | **2.5899** |
| **63** | 52-54 | 60866.89 | 3803.55 | **4061.53** | 4347.59 |  | **15** | **35.922** |
| **64** | 30-32 | 61961.48 | **2695.66** | 2817.19 | 2951.39 | 3098.75 | **23** | **10.4374** |
| **65** | 32-35 | 64675.87 | 3805.46 | **4043.26** |  |  | **16** | **0.2456** |
| **66** | 55-61 | 69131.51 | 4066.89 | **4321.84** | 4610.42 |  | **16** | **10.7086** |
| **67** | 43-48 | 70371.14 | 3910.88 | **4140.31** | 4398.98 |  | **17** | **5.6966** |
| **68** | 27-30 | 71546.5 | 3767.19 | **3976.03** | 4208.75 |  | **18** | **13.3582** |
| **69** | 43-46 | 73503.12 | 3869.51 | **4084.47** | 4324.86 |  | **18** | **2.0896** |
| **70** | 38-47 | 78573.65 | 4366.91 | **4622.2** | 4911.91 |  | **17** | **13.0042** |
| **71** | 56-60 | 82268.89 | 4114.74 | **4330.4** | 4571.76 |  | **19** | **9.0467** |
| **72** | 49-51 | 82515.95 | **5158.18** | 5502.15 |  |  | **16** | **1.6811** |
| **73** | 48-52 | 82519.99 | 4344.28 | **4585.33** |  |  | **18** | **3.0919** |
| **74** | 35-37 | 87321.2 | 4368.42 | **4597.26** | 4850.26 |  | **19** | **31.5468** |
| **75** | 30-31 | 89913.83 | 4496.35 | **4733.9** | 4995.99 |  | **19** | **9.7397** |
| **76** | 40-42 | 91354.13 | 4569.08 | **4808.2** | 5076.8 |  | **19** | **15.1919** |
| **77** | 41-46 | 95149.49 | 2213.68 | **2266.58** |  |  | **42** | **6.454** |
| **78** | 31-32 | 108555.4 | 3393.29 | **3502.86** |  |  | **31** | **2.8761** |
| **79** | 56-58 | 110410.72 | **5259.15** | 5521.03 |  |  | **21** | **14.5258** |
| **80** | 61-62 | 122446.53 | 4711.93 | **4897.37** |  |  | **25** | **52.9783** |
| **81** | 44-49 | 123343.93 | **5607.28** | 5874.81 |  |  | **22** | **9.3844** |
| **82** | 61-62 | 128785.11 | 5152.55 | **5366.91** |  |  | **24** | **4.8809** |
| **83** | 62-63 | 135746.25 | 5656.4 | **5903.75** |  |  | **23** | **23.7994** |
| **84** | 28.5-31 | 138826.22 | 5341.98 | **5552.94** | 5784.47 | 6037.45 | **25** | **31.53** |
| **85** | 35-40 | 143472.97 | 5314.7 | **5518.33** | 5740.96 |  | **26** | **24.3785** |
| **86** | 30-32 | 146238.89 | 5849.09 | **6093.95** | 6361.18 |  | **24** | **41.573** |
| **87** | 40-42 | 151213.73 | 6049.98 | **6300.58** | 6576.1 |  | **24** | **20.8207** |
| **88** | 35-38 | 162581.08 | **6022.39** | 6254.27 |  |  | **27** | **5.3015** |
| **89** | 61-62 | 162876.48 | 6265.67 | **6515** | 6788.55 |  | **25** | **24.63** |
| **90** | 42-44 | 163801.54 | 6069.35 | **6299.76** | 6552.68 |  | **26** | **39.7447** |
| **91** | 62-63 | 178543.76 | 6612.94 | **6867.72** | 7143.99 |  | **26** | **27.385** |
| **92** | 61-62 | 185058.15 | 6609.85 | **6853.92** | 7120.17 |  | **27** | **35.9757** |
| **93** | 62-63 | 185697.98 | 6633.98 | **6877.05** | 7143.99 |  | **27** | **38.925** |
| **94** | 58-60 | 190668.78 | 6335.82 | 6574.43 | **6812.89** |  | **28** | **55.8796** |
| **95** | 36-40 | 199271.74 | 6872.79 | **7117.5** |  |  | **28** | **14.0733** |
| **96** | 60-61 | 209054.06 | 6745.38 | **6968.13** | 7210.42 |  | **30** | **34.9998** |
| **97** | 60.5-62 | 318265.41 | 9091.2 | **9364.16** | 9644.95 | 9948.12 | **34** | **82.6864** |
| **98** | 60.5-62 | 339951.94 | 9185.91 | **9444.99** | 9716.18 |  | **36** | **98.6534** |
| **99** | 29-30 | 386711.47 | 8994.66 | **9209.14** | 9431.88 |  | **42** | **40.2303** |

**Note:** Bold *m/z* among each row is the most abundant (based on the most intense signal) for each protein. The *m/z* for each protein in each row is consecutive from smaller to larger, from left to right.

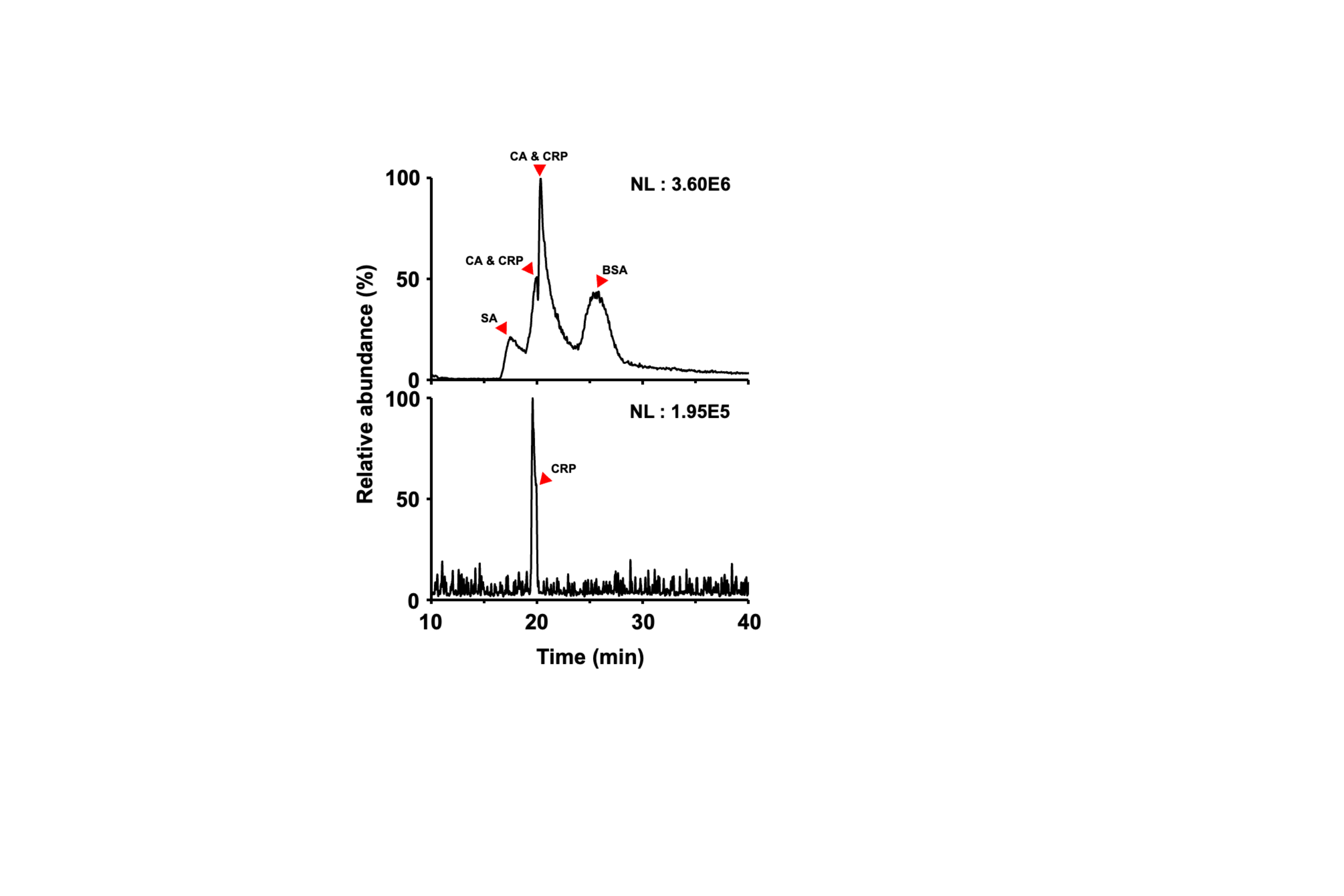

**Figure S1.** The data of a mixture of standard protein complexes analyzed by CZE-ESI-UHMR, original concentration (13 μM, top) and 50-times dilution (bottom). SA: Streptavidin; CA: carbonic anhydrase; CRP: C-reactive protein; BSA: bovine serum albumin. Cyt C, Myo, and GDH were not detected in the runs. Relative abundance labels are applied by the Excalibur software and indicate normalization of the y axis.

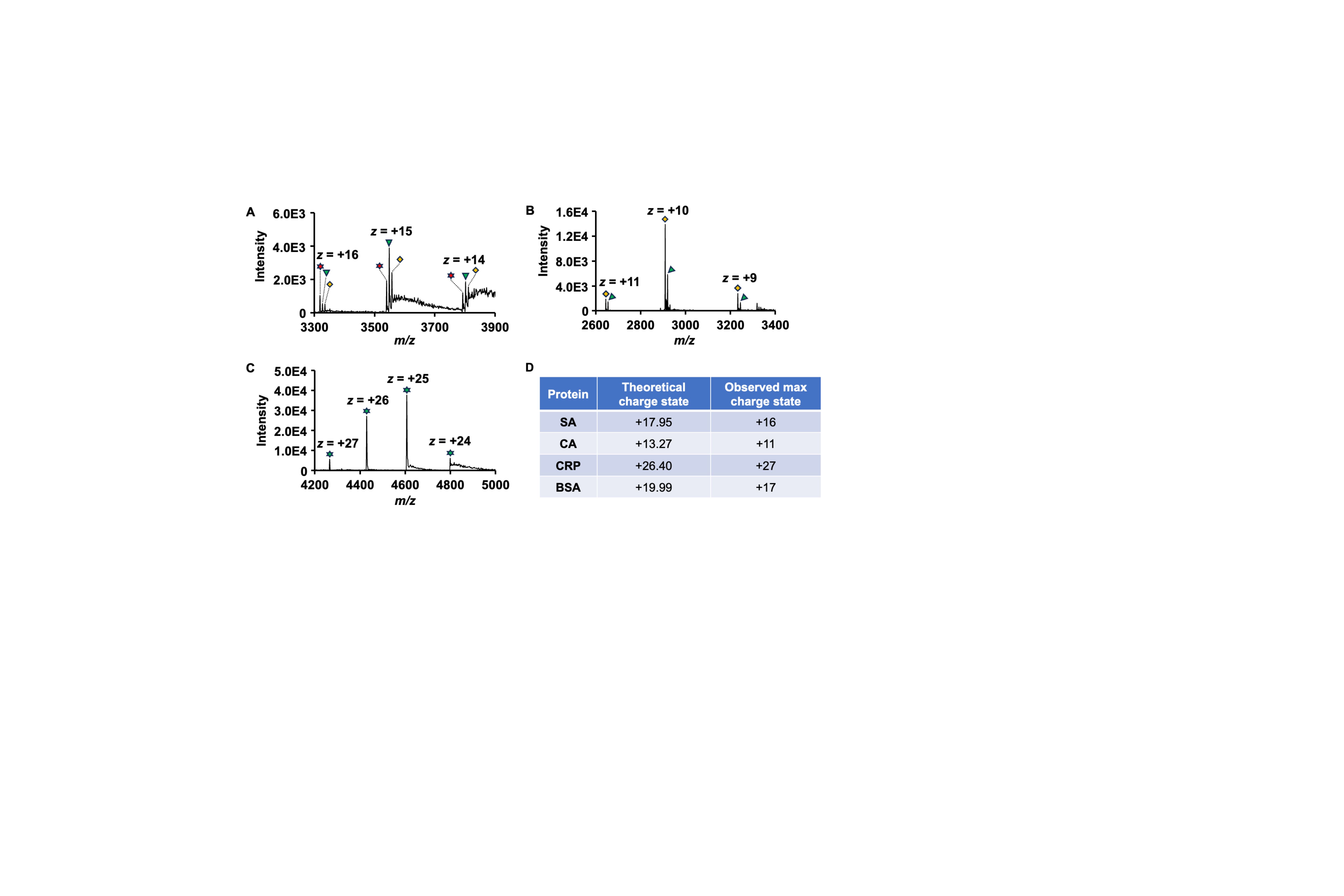

**Figure S2**. The mass spectra of some standard protein complexes from Figure S1. (A) Tetramer of SA (red star: 53084.49 Da; green inverted triangle: 53215.89 Da; yellow diamond: 53348.18 Da). (B) CA (yellow diamond: 29087.73 Da; green inverted triangle: 29194.01 Da). (C) CRP (green star: 115148.23 Da). (D) Summary of theoretical charge states (calculated by Rayleigh charge ‘Z_R_’ = 0.0778*M^0.5^) and observed maximum charge states of 4 standard proteins or protein complexes.

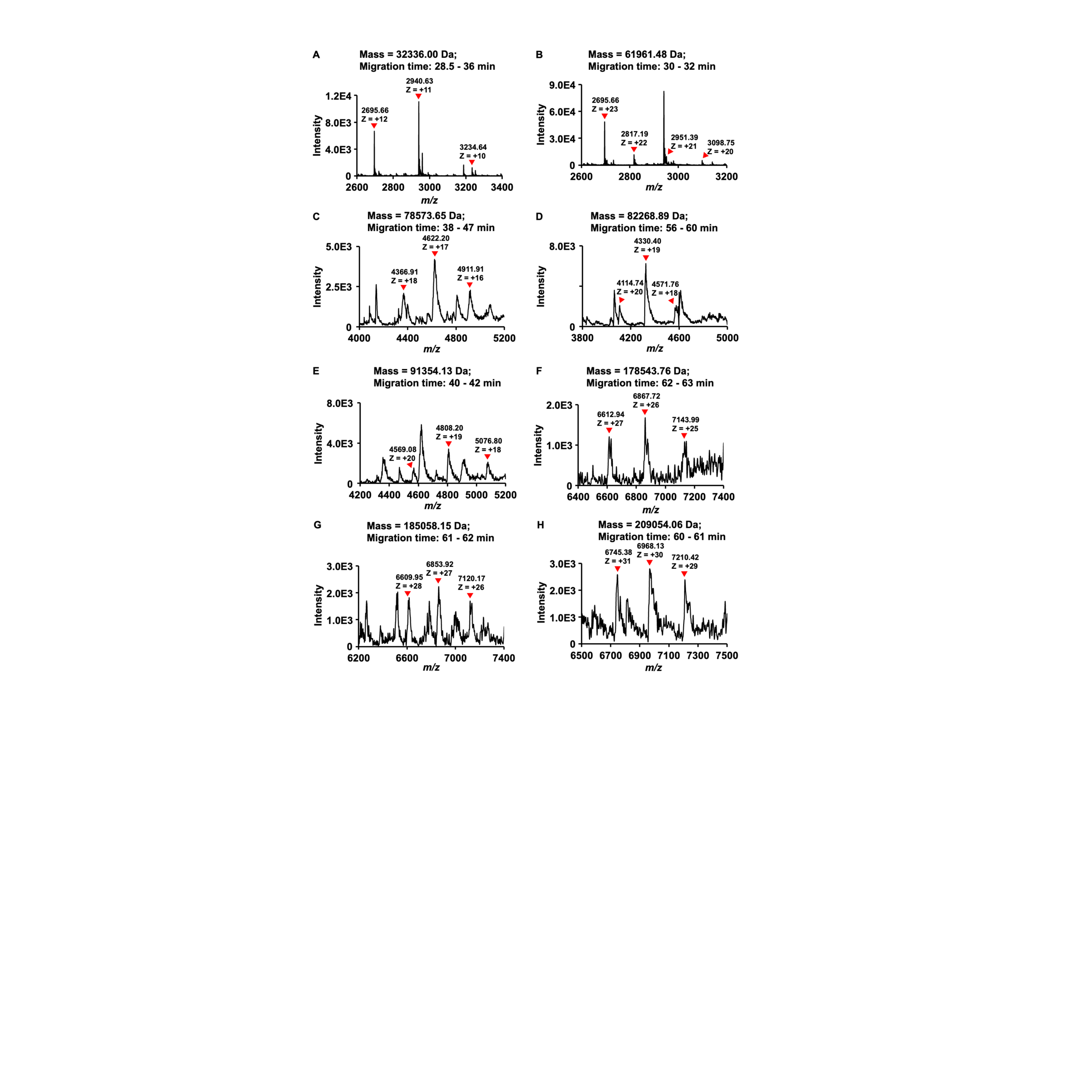

**Figure S3.** Representative mass spectra of proteoforms/protein complexes detected from the E. coli sample.

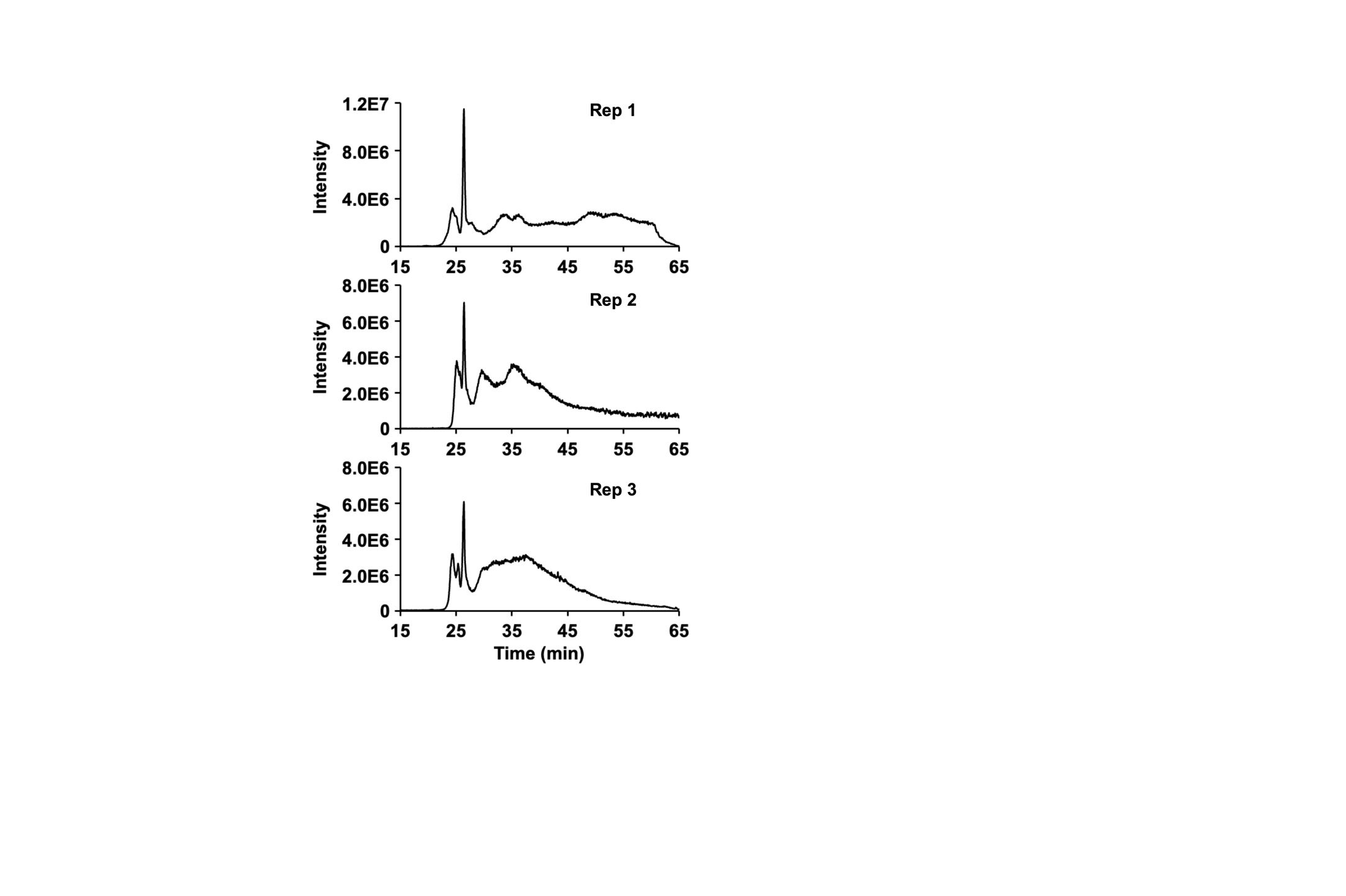

**Figure S4.** Electropherograms of triplicate analyses of an E. coli cell lysate by nCZE-ESI-UHMR. The electropherograms were aligned according to the most abundant peak.

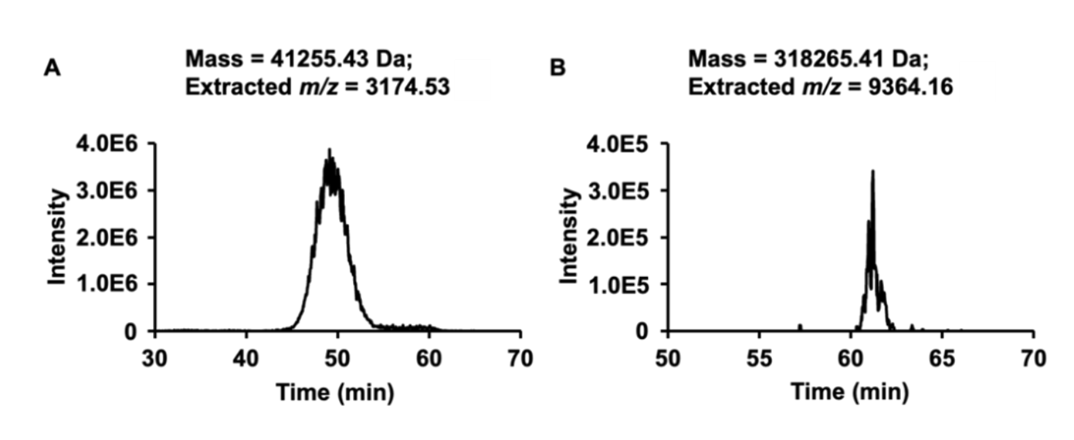

**Figure S5.** Extracted ion electropherograms of two example proteoform/protein complexes. The mass tolerance is set to 500 ppm, and Gaussian smoothing was enabled at 5 points.
